# Dynamic Electrical Stimulation Promotes hiPSC-CM Differentiation and Functionality

**DOI:** 10.1101/2024.05.01.591938

**Authors:** Nikhith Kalkunte, Sogu Sohn, Talia Delambre, Sarah Meng, Amy Brock, Janet Zoldan

## Abstract

Human induced pluripotent stem cell differentiated cardiomyocytes (hiPSC-CMs) hold great potential to resolve cardiovascular disease but are stymied by their functional immaturity. The complex electric potentials measured during cardiogenesis point to the potential of exogenous electrical stimulation in improving cardiac differentiation and functionality. Herein, we create, validate, and implement a low-cost electrical stimulation device to stimulate hiPSCs during cardiac differentiation. Notably, our open-source device enables the generation of complex electrical stimulation regimes that may vary in frequency and pulse duration over time. Our results demonstrate that dynamic stimulation during differentiation improves cardiac differentiation efficiency, calcium handling, and flow velocity, and promotes significant transcriptomic pathway enrichment compared to static stimulation or no stimulation controls. Moreover, dynamic stimulation enhances electrochemical coupling and promotes the expression of cardiogenic pathways, potentially via sarcomere development. We anticipate that more complex dynamic electrical stimulation regimens may be generated to further optimize hiPSC-CM functionality and maturity.

## Introduction

Human induced pluripotent stem cell differentiated cardiomyocytes (hiPSC-CMs) have the potential to improve the clinical outcomes of myocardial infarction due to their low immunogenicity and ability to be generated to an essentially limitless degree. Yet their translation to the clinic is greatly stymied by their functional immaturity as compared to the myocardium they seek to replace. hiPSC-CMs lack defined structure,^1,2^ generate significantly lower force during contraction,^2,3^ conduct calcium signals more slowly,^4,5^ and are metabolically inefficient^6,7^ compared to native myocardium.

Electric fields and their effect on cardiac contraction and signal propagation have been explored extensively to bridge this functional gap. Cardiomyocytes (CMs), as both signal conduits and contractile units, are particularly susceptible to electrical fields generated via exogenous stimulation. Intracellular calcium levels specifically are modulated by exogenous electric fields via voltage-gated ion channels.^8^ With calcium flux a vital element of CM contraction and signaling, exogenous electric fields have unique effects on CM transcription and function.^9^ Electric stimulation (EStim) of primary myocytes and cardiomyocytes derived from human pluripotent stem cells (hPSC-CMs) improves ultrastructural organization, calcium handling, and electrochemical coupling.^10–12^ Notably, prior work predominantly introduces exogenous EStim after the completion of cardiac differentiation in an effort to improve hiPSC-CM maturity via pacing. Indeed, dynamic frequency stimulation, accomplished via linear increases in EStim frequency over time, mimics the mechanical loading during the fetal-postnatal transition and improves tissue morphology, gene expression, calcium holding, and force-frequency response.^13^

This strategy of maturation does not leverage what we know of electric activity changes throughout embryogenesis. Assessment in animal models shows endogenous electric current rises sharply as early as Carnegie Stage (CS) 6, during gastrulation and mesoderm streak formation, and is vital to healthy cardiac development.^14–17^ In fact, EStim during cardiogenesis leads to improved outcomes. Early work in mice shows that stimulating differentiating embryonic stem cells with single monophasic pulses results in improved cardiac purity mediated via generation of reactive oxygen species.^18,19^ More recent work with hiPSCs confirms the relationship with stimulated hiPSC cultures, demonstrating improvements to cardiac purity, calcium handling, and adult-like gene expression profiles.^20–25^ This work, however, fails to capture the dynamic electric potential seen during embryogenesis. Assessments of electric charge over embryo development indicate cumulative increase over time.^14–16^

Herein, we assess the effects of dynamic EStim on cardiac differentiation and functionality by increasing frequency linearly over time. Notably, we construct and characterize a low-cost electrical stimulation device leveraging consumable electronics and provide an open-source design to support further modification by the community. Leveraging this device, we show that dynamic EStim during differentiation improves cardiac differentiation efficiency, calcium handling, and flow velocity, and promotes significant transcriptomic pathway enrichment over static electrical stimulation or no stimulation controls.

## Methods

### Device Synthesis and Validation

The low-cost electrical stimulation device is comprised of 2 parts: 1) signal generator and 2) in-well electrodes. The signal generator is assembled from an Arduino ESP32 microcontroller, L293D motor h-bridge, and LM2596 high voltage buck-buck convertor among other things (Figure 1A). The in-well electrodes are comprised of graphite carbon blocks cast in polydimethylsiloxane (PDMS). Complete fabrication instructions and program code can be found in Supplementary Data. Briefly, custom molds are 3D printed with temperature-resistant filament (Figure 1B). Carbon-graphite blocks are then cut and secured in molds (Figure 1C). PDMS (1:10 elastomer/crosslinker ratio) is poured into mold to secure carbon blocks at specific spacing for 6 well plate (Figure 1D). After PDMS curing, holes are drilled through the PMDS and into the carbon blocks to allow for electrode pin insertion (Figure 1E). Electrode pins are connected to the pulse generator. The signal generator is programmed via custom Arduino code written to generate biphasic electrical signals, measure/record voltage, current and resistance, and connect to a wireless graphic user interface (GUI). Signal generation was validated across a range of voltages and pulse durations via oscilloscope recording (Tektronix). Percent error from expected values were calculated for each measured setting. Biocompatibility of electrical stimulation was confirmed via LIVE/DEAD staining (Proteintech, PF00007) and staining for pluripotency markers after 24 and 72 hours of stimulation at tested settings (4V,5ms, 1Hz). Percent live cells was calculated by dividing the number of cells positively stained for Calcein AM (live) by the total cell number, calculated as the sum of positively stained for Calcein AM (live) and ethidium homodimer (dead).

**FIG. 1.**
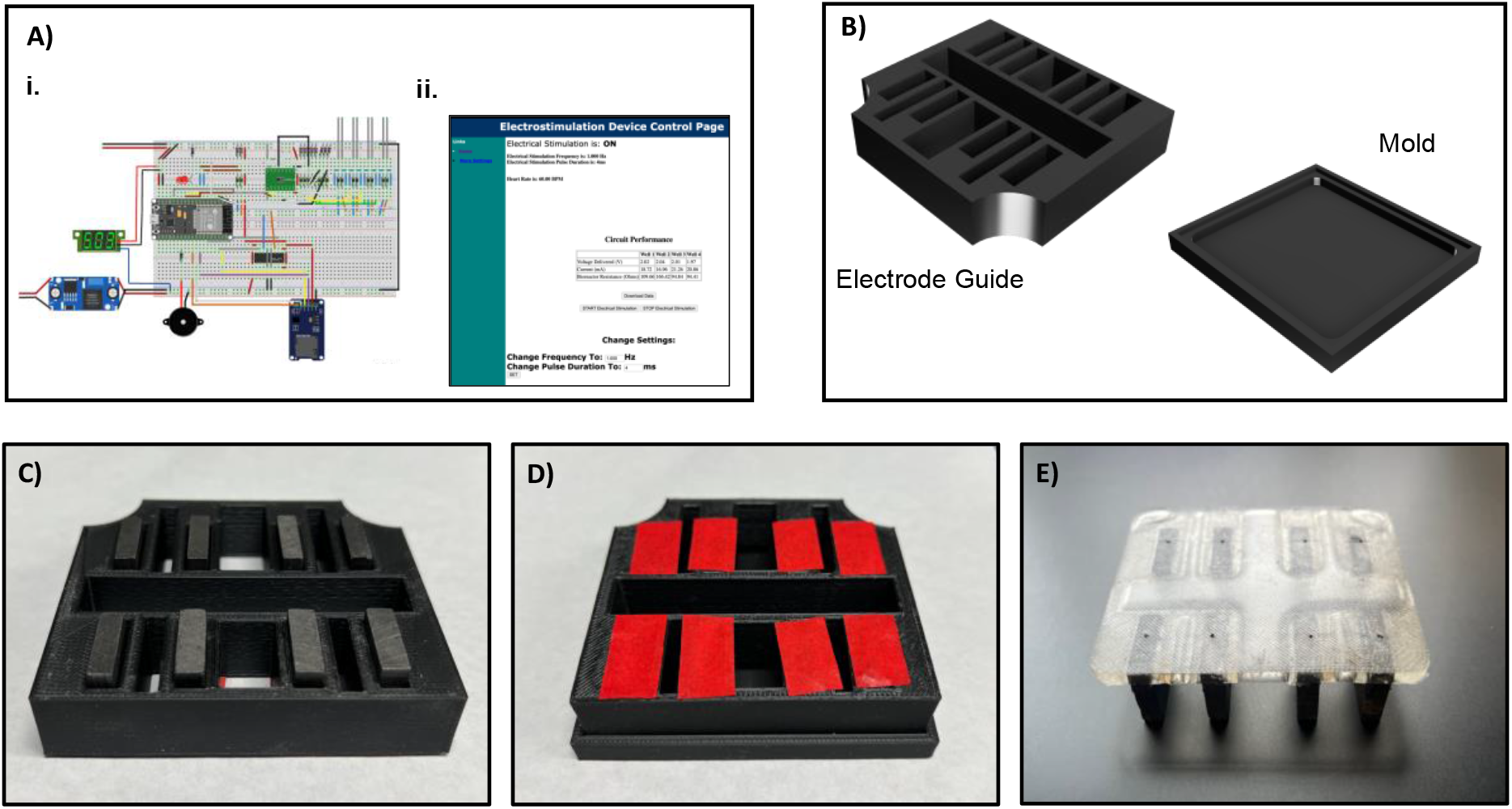
Electrical stimulation device design and manufacturing **(A)** i. Electrical stimulation voltage-sensing circuit. ii. Web server which manages stimulation parameters and enables downloads of stimulation data. **(B)** Two-part electrode building mold, with electrode guide and base mold. **(C)** Electrodes loaded into electrode guide. **(D)** Mold with PDMS poured and electrodes secured into guide with tape. **(E)** Final electrodes with wire holes drilled.

### Cardiac Differentiation and Electrical Stimulation

WTC-11 hiPSCs, transfected to express CMV-GCaMP2, a gift from Dr. Bruce Conklin, were seeded in Matrigel coated plates (0.1mg/ml) at 28,000 hiPSCs per cm^2^. Cardiac differentiation followed a modified protocol of Wnt pathway modulation.^26^ Cells were cultured in Essential 8 media for 72 hours, with media changes every 24 hours. On Day 0 of cardiac differentiation, hiPSCs were initiated with RPMI 1640 media with B27 no insulin (RB-) and 12μM CHIR99021. After 24hrs, cells were transitioned to RB-media for forty-eight hours. On Day 3 of differentiation, Wnt signaling was inhibited by incubating cells in RB-media with 5mM IWP2 (inhibitor of WNT production). Forty-eight hours after inhibition with IWP2, cells were allowed to recover in RB- for another forty-eight hours. After, cells were then fed with CDM3 media (RPMI 1640 with h-albumin [100 mg/mL] and L-ascorbic acid 2-phosphate [100 mg/mL]). Electrical stimulation was also initiated on Day 0 of cardiac differentiation. Cells were stimulated continuously from Day 0 to Day 10 of cardiac differentiation along 3 conditions (Voltage/Pulse Duration/Frequency): 4V/4ms/1Hz, 4V/4ms/2Hz, and 4V/4ms/dynamic Hz. Dynamic frequency stimulation, termed frequency ramped stimulation, followed a linear increase in frequency from 1 Hz-2 Hz at 6.94E-5 Hz/minute (equivalent to 0.1 Hz per day).

### Immunostaining

Cells differentiated under EStim and no stimulation controls were fixed and stained on Day 10 to assess functionality and cardiac purity. Cells were fixed with 4% paraformaldehyde (Polysciences) for 10 min. at room temperature (R/T), rinsed with 300 mM glycine (Sigma-Aldrich), and permeabilized with 0.2% Triton X-100 (Amresco). Cells were then incubated in blocking buffer, containing 1% BSA and 0.1% Tween-20 (Fisher Scientific), for 30 min. and incubated overnight at 4°C with primary antibody in blocking buffer. Primary antibodies, including cardiac troponin T (cTnT; 1:500, Proteintech, 15513-1-AP) and Octamer-binding transcription factor 4 (OCT-4; 1:40, Invitrogen, MA5-14845), were incubated for 24 hours. After, cells were washed with blocking buffer and incubated with secondary antibodies in blocking buffer. Secondaries used include anti-rabbit Alexa-488 (1:400, Sigma Aldrich, SAB4600044). Cells were co-stained with Draq5 (Cell Signal, 4048L) or nuclei stain 4’,6-Diamidine-2’-phenylindole dihydrochloride (DAPI; Thermo Fisher, D1306). Stained samples were imaged using an Olympus FV3000 Confocal Laser Scanning Microscope.

### Intracellular Calcium Transients Imaging

The flow of intracellular calcium was facilitated via high-speed imaging of CMs differentiated from CMV-GCaMP2 transfected hiPSCs. Briefly, on Day 10 of cardiac differentiation, GCaMP cells were imaged on a Leica DMI6000 B microscope and a 10X Leica objective at 20 frames per second. The cyclic intensity of green fluorescent protein (GFP), representing the intracellular concentration of calcium, was captured in image sequences of 120 images. Images were processed using a custom MALTAB code to assess degree of synchronicity, calcium cycle duration, and decay constant, as previously discussed.^27^ Images were also processed with the Optical Flow Analysis Toolbox in MATLAB for Mesoscale brain activity (OFAMM) to derive calcium flow directionality and velocity.^28^ Briefly, image sequences were pre-processed using median spatial filters (3X3 pixels) and low pass (<1Hz) temporal filters as recommended. Processed image sequences were then passed to the toolbox, wherein the Horn-Schunck (HS) optical flow method was employed to calculate vectors of flow on a pixel-by-pixel basis. Vector directionality was assessed via calculating circular variance of vector phase whereas velocity was calculated as vector magnitude.^29^

### Quantitative reverse transcription–polymerase chain reaction

RNA from cells on Day 10 of cardiac differentiation was isolated, reverse-transcribed, and quantified according to manufacturer’s protocols. Briefly, the RNeasy Mini Kit (Qiagen) was implemented according to manufacturer’s protocol. RNA was eluted using nuclease-free (NF) water and kept on ice during RNA quantification, which was performed using a Take3 plate in a Cytation 3 (BioTek Instruments). RNA purity was assessed (260/280 absorbance >2) and RNA samples were then kept on ice while proceeding to reverse transcription. Reverse transcription was performed according to manufacturer’s protocols via a High-Capacity cDNA Reverse Transcription Kit (Applied Biosystems). Quantitative PCR was performed using PrimeTime qPCR Primers (Integrated DNA Technologies) and PowerUp SYBR Green (Thermo Fisher) on a StepOnePlus Real-Time PCR System (Applied Biosystems) based on previously described methods.^27^ TATA-binding protein (TBP) was used as the endogenous control, and mRNA from CMs not subjected to stimulation was used as the reference sample. The full list of primers used is available in supplementary table S1.

### Gene Expression Profiling by 3’ Tag-Seq

For each condition, total RNA was collected from two technical replicates of two separate differentiations each by the RNeasy Mini Kit (Qiagen) according to the manufacturer’s protocol. Samples were submitted to the University of Texas Genomic Sequencing and Analysis Facility (Center for Biomedical Research Support, RRID: SCR 021713). Tagseq libraries were sequenced using the NovaSeq 6000 SR100. Averaging the counts across technical replicates yielded 2 samples for each condition. Reads were preprocessed by the Bioinformatics Consulting Group at the University of Texas at Austin, and reads were processed and mapped using the RNAseq pipeline by nf-core.^30^ Briefly, the reads were aligned to the GRCh38 human reference genome using ‘STAR’, and gene counts were obtained using ‘Salmon’.^31,32^

Differentially expressed genes were calculated using DESEQ2 with the adaptive shrinkage estimator, ‘ashr’.^33,34^ Differentially expressed genes were defined to have a p-adjusted value of 0.05 and a log fold change cut-off of 1. Using the differentially expressed genes, gene set enrichment analysis was performed using the fgsea package in R and the Gene Ontology Biological Processes pathway set.^35–37^

### Statistical analysis

Statistical analysis was conducted employing one-way analysis of variance tests followed by post hoc Tukey’s multiple comparison tests using Prism 9 software (GraphPad). Relationships were deemed significant at a threshold of p<0.05. The data are presented as mean values ± standard error. Statistical tests were performed after Robust regression and Outlier removal (ROUT) at Q=1% (GraphPad).

## Results

### Device Characterization

Oscilloscope recordings of generated signals confirm the generation of biphasic, monophasic, and alternating monophasic pulses (Figure 2A, Supplementary Data S1). Signal fidelity assessments, calculated as percent error between measured and expected signal parameters, yields <5% across all durations at pulse durations greater than 2ms, with 5ms being an exception resulting in 20% error (Figure 2B). Electrical stimulation of hiPSCs at a range of pulse durations and frequencies over 72 hours, resulted in non-significant differences in %live cells as compared to no stimulation controls (Figure 2C). Immunostaining of hiPSCs, post 72 hours of stimulation at 1Hz, 4ms, 4V, shows colocation of OCT-4 transcription factor with DAPI nuclear stain, indicating retention of pluripotency (Figure 2D). Voltage sensing modules show high accuracy in sensing voltage and current (<10% error), while sensed resistance is prone to sensor noise (Supplementary Data S2). Measurement of frequency over 10 days of stimulation in frequency ramped stimulation show linear increases in stimulation frequency at 0.1 Hz per day rate (Supplementary Data S3).

**FIG 2.**
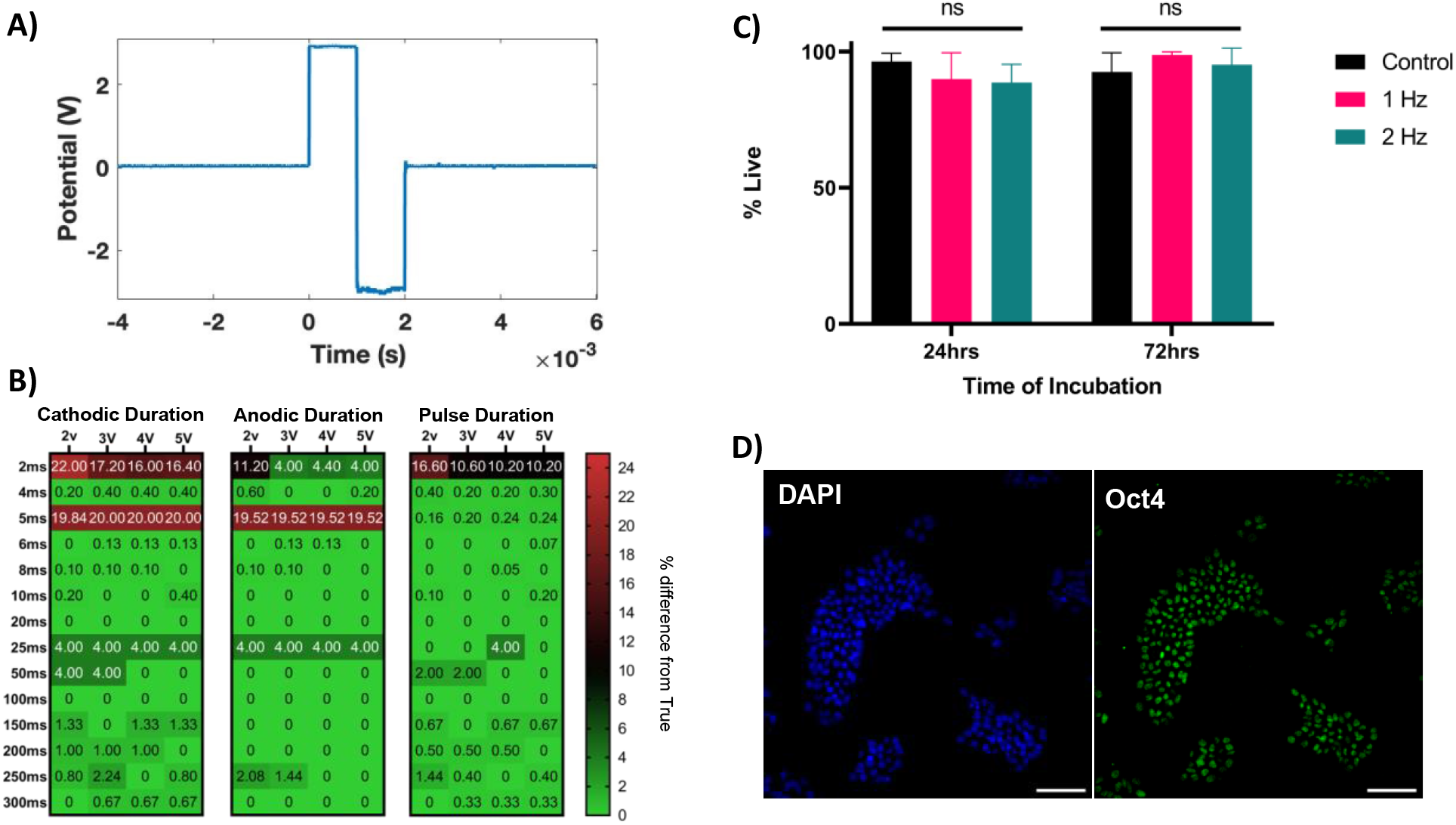
Electrical stimulation signal assessed for signal fidelity and biocompatibility. **(A)** Oscilloscope recordings of recorded signals demonstrate generation of biphasic, monophasic, or alternating monophasic signal pulses. **(B)** Signal fidelity of biphasic stimulation is assessed via cathodic duration (duration of positive voltage), anodic duration (duration of negative voltage) and total pulse duration. Percent error or measured pulse parameters compared to expected are assessed across voltages and pulse durations. **(C)** Minimal cell death is seen in hiPSCs when subjected to biphasic stimulation (4V,4ms) at 1Hz or 2Hz over the course of 72 hours (n=6). **(D)** Representative images of hiPSCs after 72 hours of electrical stimulation (1hz, 4v,4ms) show nucleus colocation of pluripotency marker Oct4. Scale bar=100μm).

### Cardiac Differentiation

Electrical stimulation during cardiac differentiation, across all regimens, is shown to significantly increase the percentage of differentiated cardiomyocytes in heterogeneous cell cultures. Firstly, electrical stimulation does not inhibit the generation of functional CMs, as indicated by spontaneous beating and immunostained sarcomeres (Figure 3A). Next, electrical stimulation at 1 Hz (0.47±0.05), 2 Hz (0.46±0.05), and ramped frequency (0.54±0.05) all result in significantly higher normalized cTnT area as compared to no stimulation controls (0.29±0.03). Non-significant differences exist between stimulation regimens (Figure 3B). Finally, cardiac subtype analysis of Day 10 hiPSC-CMs reveals electrical stimulation impacts the specialization of CMs into ventricular cardiac tissues. Expression of ventricular cardiac marker *IRX4* is upregulated to a statistically insignificant degree in CMs differentiated under 2Hz and ramped frequency stimulation, while 1Hz stimulation triggers a 5-fold downregulation, relative to control. Similarly, atrial cardiac marker *NR2F2* is downregulated 15-fold in CMs differentiated under 1Hz, 25-fold under 2Hz, and 30-fold fold in CMs differentiated under ramped frequency stimulation, relative to control (Figure 3C).

**FIG 3.**
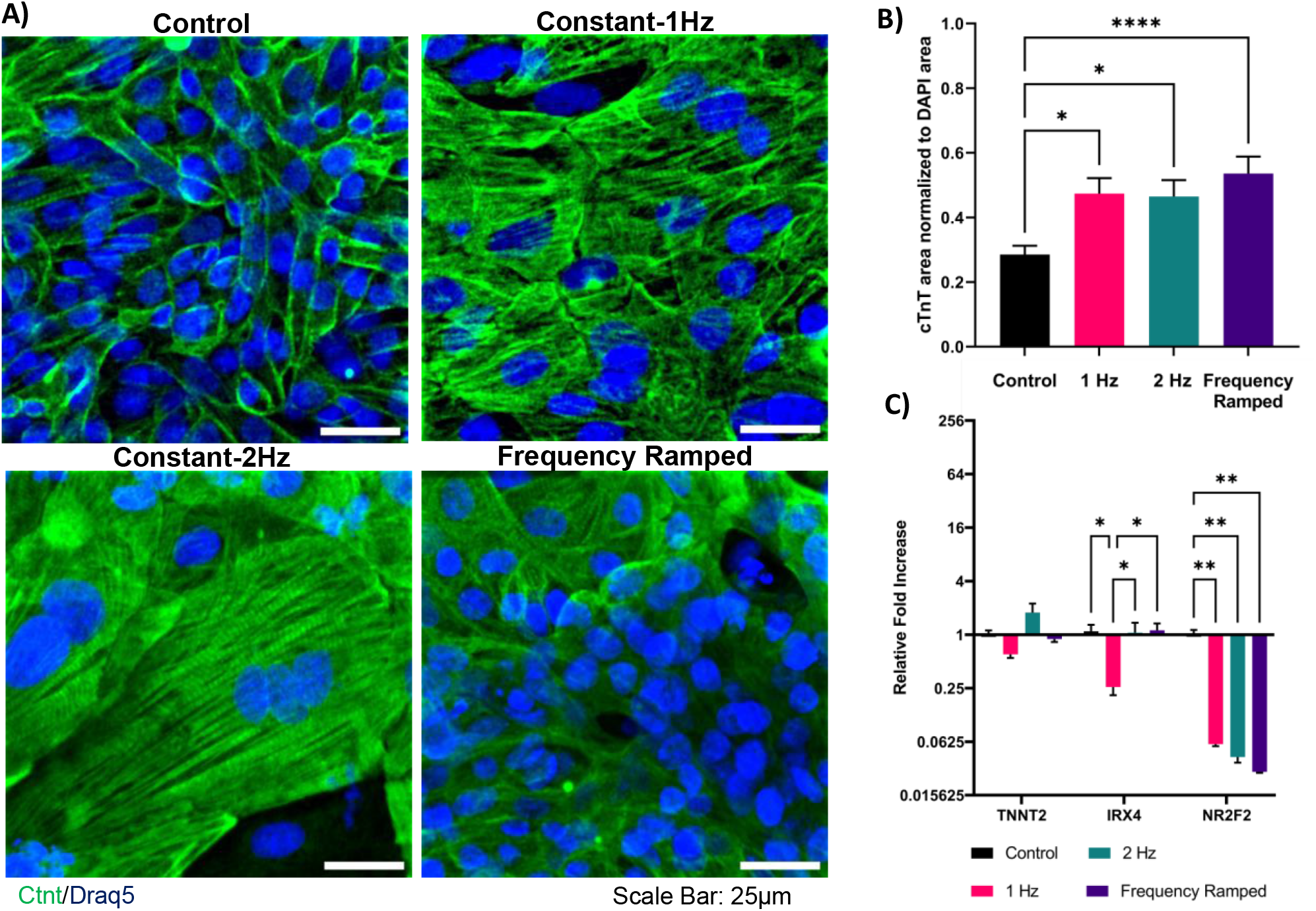
hiPSCs are differentiated under electrical stimulation and are assessed for cardiac purity. **(A)** hiPSC-CMs differentiated under electrical stimulation are stained for cTnT(green) and nuclei (blue). Sarcomeres are present in all conditions. Scale bar = 25μm. **(B)** hiPSC-CMs differentiated under electrical stimulation are assessed for cTnT expressions via immunostaining. cTnT area is normalized to DAPI area (n=20). **(C)** hiPSC-CMs differentiated under electrical stimulation are assessed for mRNA expression of canonical cardiac subtype genes (n=6). *p<0.05, **p<0.01.

### Electrochemical coupling

Assessment of intracellular calcium transients demonstrates significantly different calcium handling when hiPSC-CMs are differentiated under electrical stimulation, particularly at 1 Hz. Stimulation at 1 Hz (134.20±4.53) is shown to significantly decrease the degree of synchronicity as compared to 2 Hz (105.00±3.00), frequency ramped (97.36±2.87), and no stimulation controls (97.77±2.53). Electrical stimulation at 1 Hz also shortens the duration of calcium cycles as compared to 2 Hz, ramped frequency, and no stimulation controls. The full-width half-max of intracellular calcium transients after stimulation at 1Hz (5.31±0.18ms) are significantly smaller than those measured in the 2 Hz (9.65±0.55ms), ramped frequency (6.92±0.23ms) and no stimulation controls (7.42±0.15ms). Notably, 2 Hz stimulation drives significantly longer calcium cycles as compared to control and frequency-ramped conditions (Figure 4). Electrical stimulation also influences the uptake rate of calcium. The calcium cycle decay constant (τ), a measure of intracellular calcium reuptake, is seen to be significantly higher in 1 Hz (12.51±0.41ms) stimulated cells compared to 2 Hz (9.99±0.0.27ms), frequency ramped (9.73±0.30ms), and no stimulation controls (11.22±0.34ms). Additionally, CMs from the 2 Hz and frequency ramped conditions show faster calcium reuptake when compared to no stimulation controls, as demonstrated by a significantly smaller τ (Figure 4). Gene expression analysis of critical calcium handling proteins show significant differences in expression across stimulation condition. Calcium release gene *RYR2* is significantly upregulated in CMs under 1 Hz stimulation (31.62±2.01) compared to 2Hz (11.72±2.07) and ramped frequency (16.15±0.58) conditions. Significant upregulation of connexin-43 gene *GJA1* is seen across all electrical stimulation regimens as compared to control, with 2Hz triggering the most significant upregulation (35.63±3.22) as compared to 1Hz (22.03±0.77) and ramped frequency (11.01±0.24). Significant, but marginal upregulation in *CACNA1C* (encoding for the alpha1 subunit of voltage-dependent calcium channel) is seen across all electrical stimulation regimens, relative to no stimulation (Supplementary Data S4).

**FIG. 4.**
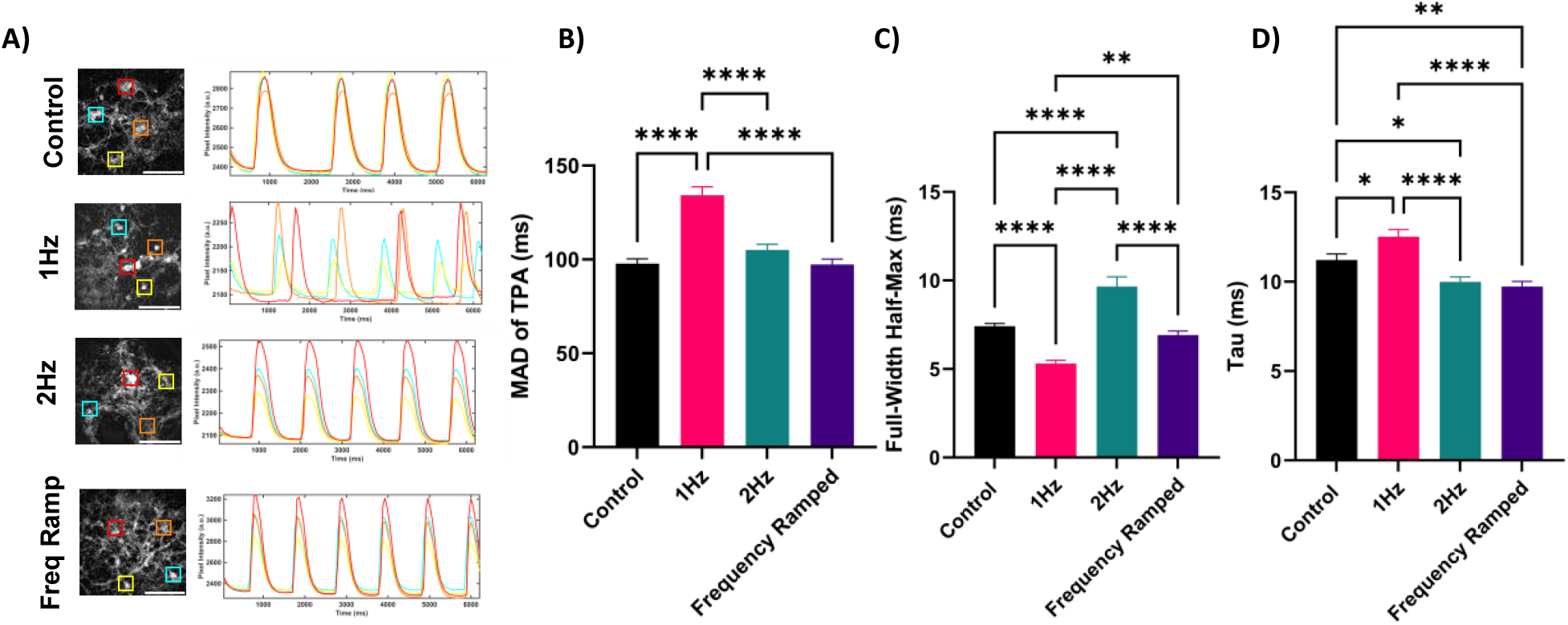
Intracellular calcium handling and synchronicity improve with dynamic frequency stimulation. **(A)** Representative calcium traces of GCaMP hiPSC-CMs differentiated under electrical stimulation. Scale bar = 500μm. **(B)** Synchronicity (n=215), **(C)** calcium cycle duration (n=175), and **(D)** decay constant (n=175) after 10 days of stimulation. *p<0.05,**p<0.01, ****p< <0.0001. TPA-MAD, time of peak arrival median absolute deviation.

### Calcium Flow Velocity and directionality

Analysis of calcium transient flow illustrates an effect of electrical stimulation on calcium flow velocity and directionality. Constant stimulation, at 1Hz (1.97±0.20 μm/s) and 2Hz (3.42 ± 0.27 μm/s) significantly decreases the velocity of calcium flow as compared to no stimulation controls (5.16±0.36 μm/s) and ramped frequency stimulation (6.13±0.52 μm/s) conditions. Ramped stimulation results in a statistically non-significant increase in calcium flow velocity as compared to no stimulation controls (Figure 5E). Ramped electrical stimulation also looks to improve the directionality of calcium flow. Assessed via variance in flow vector phase, ramped stimulation drives the lowest phase variance (0.12±0.01) as compared to control (0.22±0.02), 1Hz (0.49±0.04), and 2Hz (0.58±0.03). We see constant stimulation, at 1Hz and 2Hz result in higher variance in calcium vector phase, indicating more multidirectional calcium flow/propagation within differentiated cardiac syncytium (Figure 5F).

**FIG 5.**
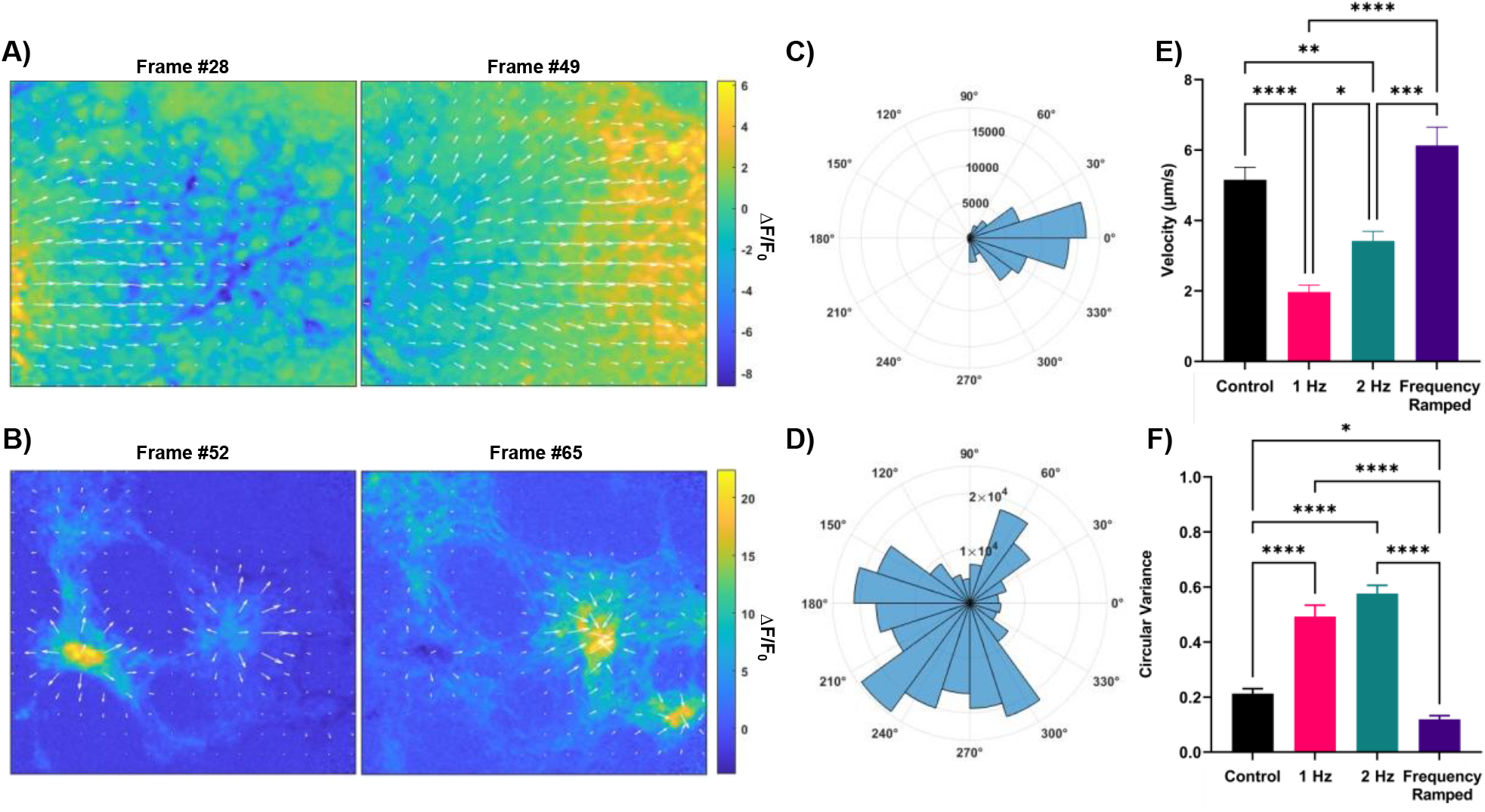
Calcium propagation velocity and directionality improve with dynamic electrical stimulation. Representative optical flow-derived vector plots of unidirectional **(A)** and multidirectional **(B)** calcium flow over the course of cardiac contraction. Representative polar rose plots display vector phase variance in unidirectional **(C)** and multidirectional **(D)** vector fields. **(E)** Average vector magnitude of calcium flow is calculated across electrical stimulation condition (n=40). **(F)** Average circular variance of vector phase is calculated across electrical stimulation condition (n=45). ΔF/F0. Change in fluorescence normalized to baseline fluorescence. *p<0.05, **p<0.01, ***p<0.001, ****p<0.0001.

### RNA sequencing and pathway expression

We performed Tag-seq analysis to characterize the effect of electrical stimulation on the transcriptome of differentiated cells. Day 10 hiPSC-CMs subjected to ramped stimulation was compared to 1 Hz constant stimulation, and no stimulation controls. Principal component analysis of all samples results in clustering within each stimulation condition, as shown by k-means clustering, demonstrating distinct transcriptomic profiles between groups (Supplementary Data S5).

We conducted differential gene analysis and pathway enrichment analysis to elucidate any condition-specific signatures. Comparing electrical stimulation conditions to non-stimulation controls highlight regimen-specific effects. A constant stimulation regimen leads to significant downregulation across the transcriptome. Control samples see an enrichment of cardiac muscle development pathways such as cardiac muscle contraction and cardiac myofibril assembly. The control group is also consistently upregulated in pathways such as chromosome segregation and organization, and cell division, showing that there is a change in cell cycle regulation due to constant electric stimulation (Figure 6A). Comparing ramped stimulation to controls yields no significant pathway enrichment. Finally, between 1Hz and ramp, the ramped stimulation method results in upregulation of pathways relating to heart morphogenesis, cardiac muscle cell differentiation, and cardiac cell development indicating more efficient development of specialized structural and/or functional features of cardiac cells due to ramped electric stimulation. The ramped condition is also upregulated in pathways such as cardiac conduction system development, regulation of cardiac muscle contraction, and regulation of heart contraction, suggesting that the ramped condition yields cells that are more functionally mature than cells exposed to constant stimulation (Figure 6B).

**FIG 6.**
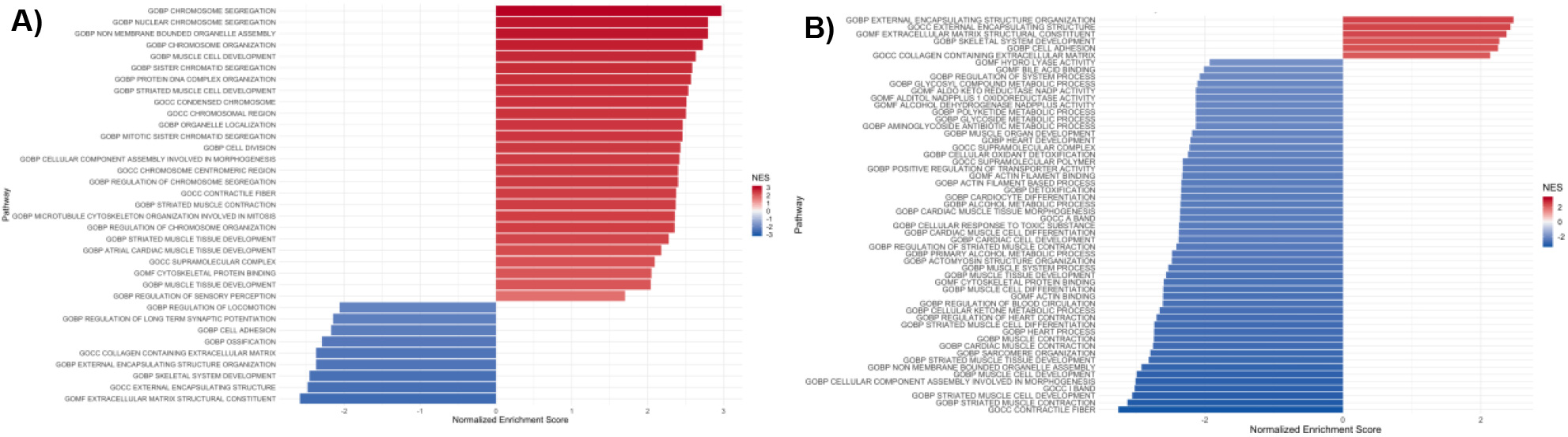
Bar plot of the results from a gene set enrichment analysis of **(A)** 1Hz vs control and **(B)** 1Hz vs frequency ramped. Pathways shown are significant (p-value < 0.05) and are colored by normalized enrichment score (NES).

To investigate the specific genes that may be driving pathway enrichment trends, we resolved the most differentially expressed genes across cardiac-specific pathways. Using pheatmap, the most differentially expressed genes across condition were plotted in a heatmap and normalized across rows; values indicated z-score within each row.^38^ The top five most differentially expressed genes include *ANKRD1, SLC8A1, MYH6, MYH7*, and *TTN* (Supplementary Data S6). Except for *SLC8A1*, all other genes relate directly to contractile protein machinery, more specifically sarcomere development and stabilization. This indicates that ramped frequency may specifically improve hiPSC-CM functionality via sarcomere development.

## Discussion

Herein, we investigate the effect of dynamic electrical stimulation on hiPSC-CM differentiation, electrochemical coupling, and transcriptome expression. We show that dynamic electrical stimulation improves cardiac differentiation efficiency, improves calcium handling and electrochemical coupling, and promotes the expression of cardiogenic pathways as compared to static electrical stimulation and no stimulation controls.

Prior work independently examines the effect of constant electrical stimulation during differentiation^20^ or dynamic stimulation post differentiation,^13^ but fails to recapitulate dynamic electrical current changes measured during embryogenesis. Our approach, including the development of an open-source, low-cost electrical stimulation system that allows for dynamic frequency stimulation, allows cells to be subjected to electrical stimulation during differentiation. Our open-source instrumentation allows for input of custom stimulation regimens varying both stimulation frequency and pulse duration over time and promotes future innovation and augmentation of electrical stimulation systems.

Our results on the promotion of cardiac differentiation via electric stimulation are well supported. Numerous stimulation regimens performed on differentiating PSCs result in increased CM purity and spontaneously beating clusters.^18–20,25^ We add to this body of research the effects of dynamic frequency EStim on cardiac differentiation.

Notably, our findings on the underperformance of 1Hz stimulated cells, relative to no stimulation control cells, differ from other studies. Several factors may contribute to this difference. First is variance in stimulation protocols. On deeper investigation, we note other studies that stimulate at 1Hz only stimulate for portions of each day of differentiation,^20,21,24,25^ whereas we stimulate continuously throughout cardiac differentiation. This is to provide a more robust comparison point to dynamic frequency EStim conditions that also vary continuously. Next, could be the physiologic irrelevance of 1Hz stimulation during cardiogenesis with reference to higher frequency and dynamic frequency EStim regimens. Fetal heart rates are well documented to range from 110-170 beats per minute,^39,40^ more closely aligning with higher frequency stimulation. This higher frequency constant stimulation is found to have positive effects on cardiac development and function.^23^ A daily analysis of differentiating cells could reveal at what point during the differentiation process the 1Hz pacing starts to negatively impact cardiac functionality. Finally, analysis timepoints may explain differences in CM functionality. We notably assess CM functionality at the termination of EStim (Day 10 of cardiac differentiation) to specifically assess the effects of electrical stimulation on CM differentiation and transcription. This timepoint aligns with Carnegie stage 8-9,^41^ still early in embryonic development. Analysis of CM functionality at alternative timepoints, may reveal more interesting long-term effects of EStim on CM functionality.

In summary, our investigation shows that differentiating hiPSC-CMs under electrical stimulation improves differentiation yield and that dynamic frequency promotes improvements in cardiac transcription and functionality. More specifically, we find that dynamic frequency EStim improves electrochemical coupling, calcium flow velocity, and directionality as compared with constant 1 Hz EStim. In addition, we release an open-source, low-cost electrical system to the wider community to increase access to this work and promote technical innovation and experimental complexity. We see this platform, along with our initial analysis of dynamic frequency stimulation, as a means to spur further research into the role electrical stimulation may play in hiPSC-CM functionality and maturation.

