## Supplementary Data for "Dynamic Electrical Stimulation Promotes hiPSC-CM Differentiation and Functionality"

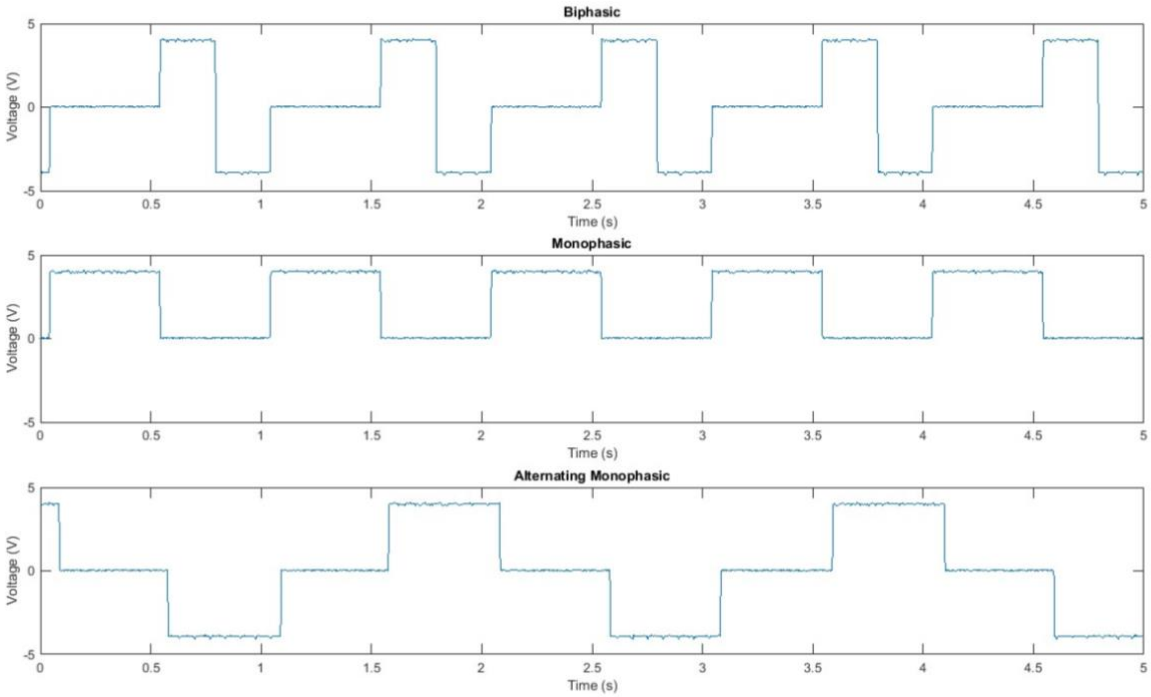


**S1: Pulse generation:** Plotting collected oscilloscope data of signal generation show successful generation of biphasic, monophasic, and alternative polarity monophasic signals.

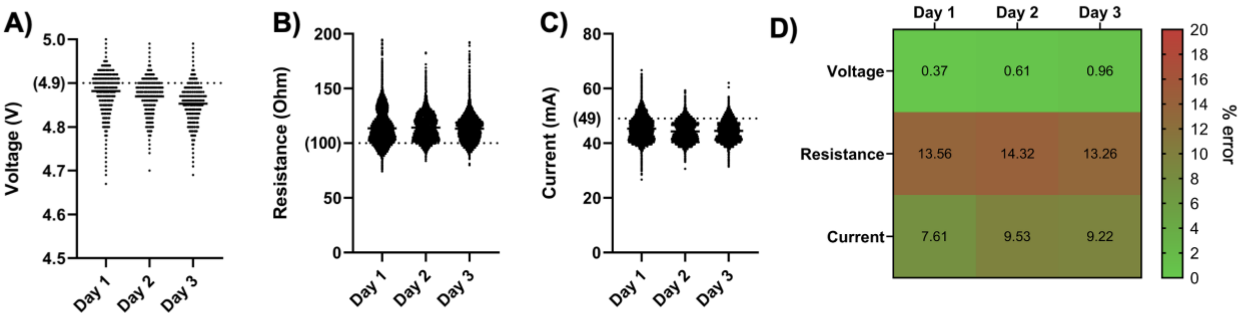


**S2: Pulse Sensing Performance:** Plotting measured voltage, resistance, and current over 72 hours show little variance. Voltage **(A)** shows less than 1% error in measured vs actual, indicated by the dotted line. Measured resistance **(B)** and current **(C)** exhibit a 10% and 15% error when compared to expected values **(D)**.

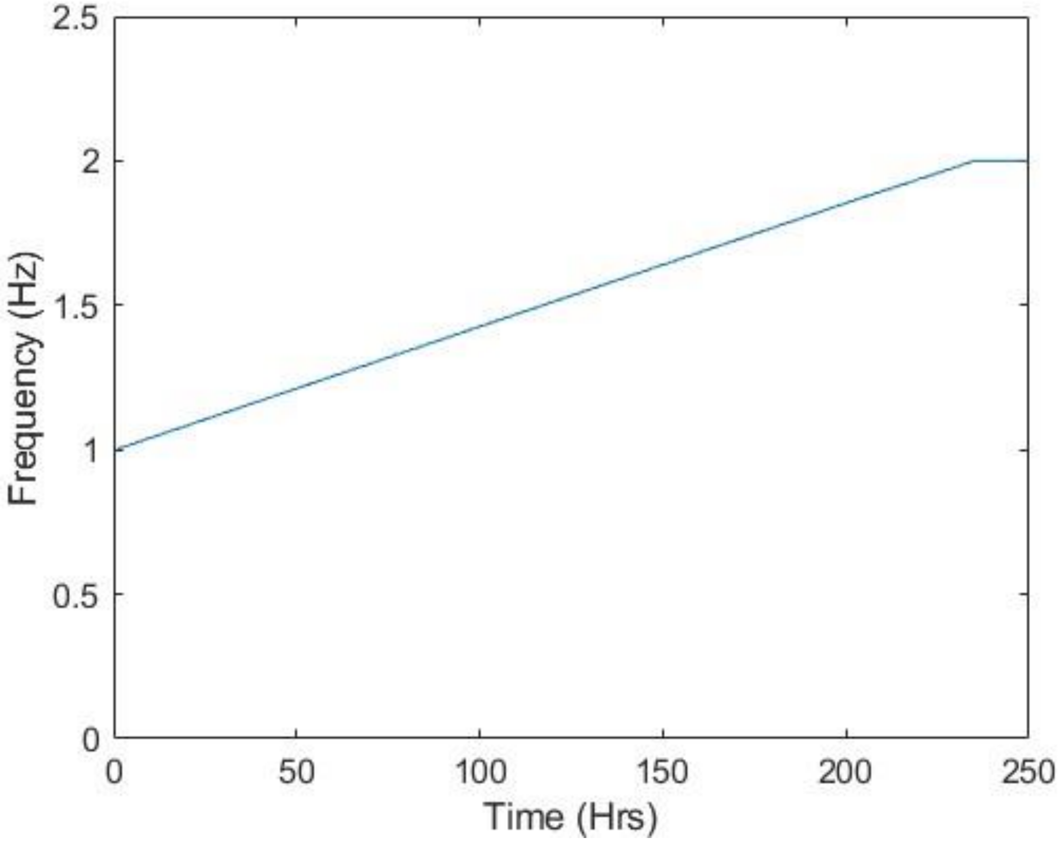


**S3: Dynamic Frequency:** Tracking the frequency of electrical stimulation over 10 days of stimulation for Ramped Frequency conditions show a linear increase in frequency form 1Hz to 2HZ, as programmed. Through user defined stimulation parameters we achieve an increase rate of 0.0043 Hz/hr (0.1Hz/day), as desired.

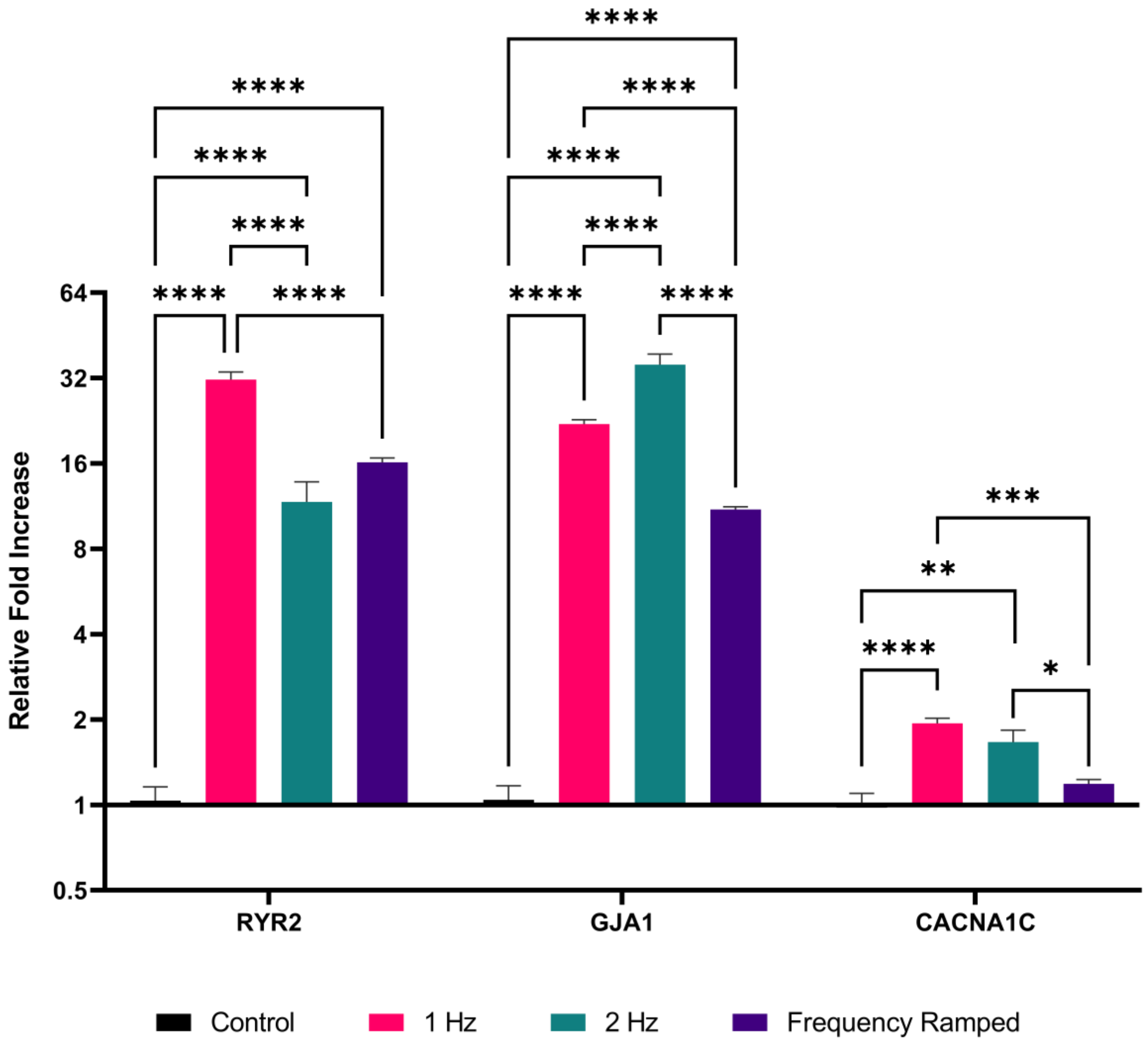


**S4: Calcium Handling Gene Expression:** hiPSC-CMs differentiated under no stimulation (Control), 1 Hz, 2Hz, and frequency ramping are assessed for mRNA expression of calcium handling genes at day 10 relative to control (n=6). *p<0.05, **p<0.01, ***p<0.001, ****p<0.0001.

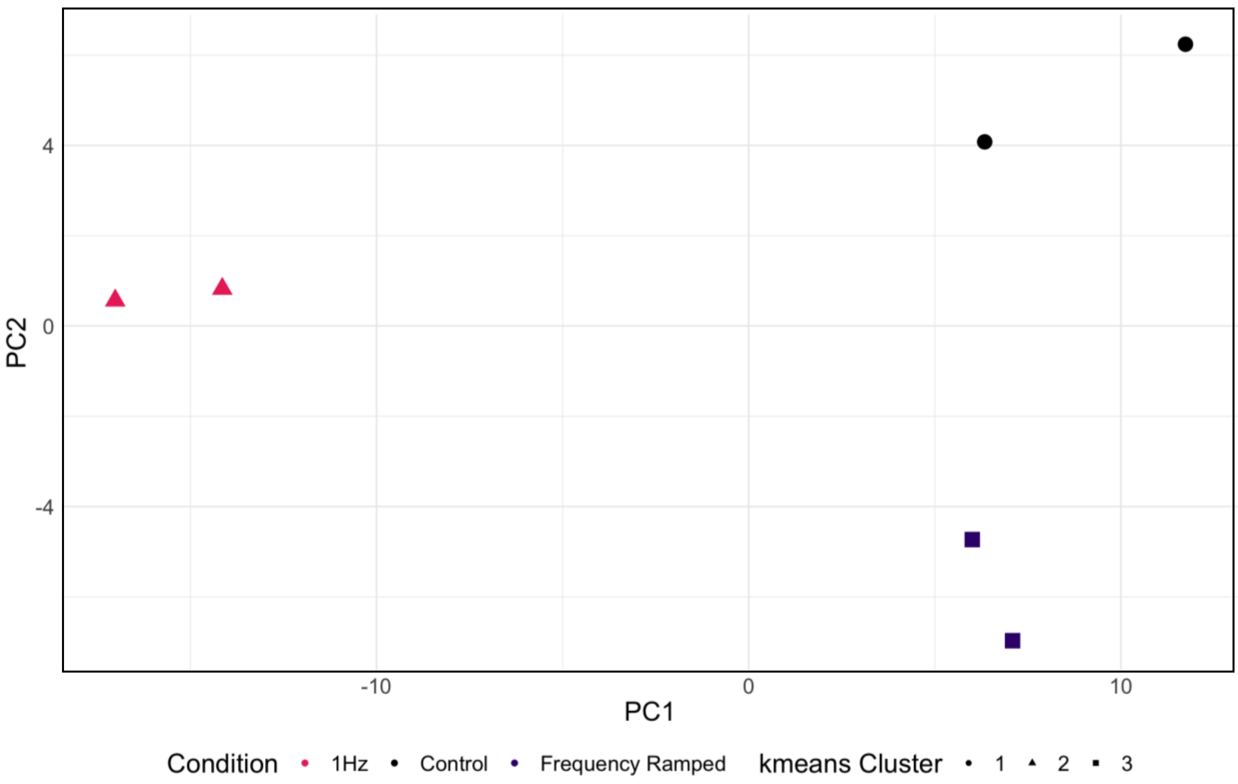


**S5: PCA of RNA-Seq Data:** hiPSC-CMs differentiated under no stimulation (Control), 1 Hz, 2Hz, and frequency ramping are processed via RNA sequency and assessed for clustering via principal component analysis and kmeans clustering.

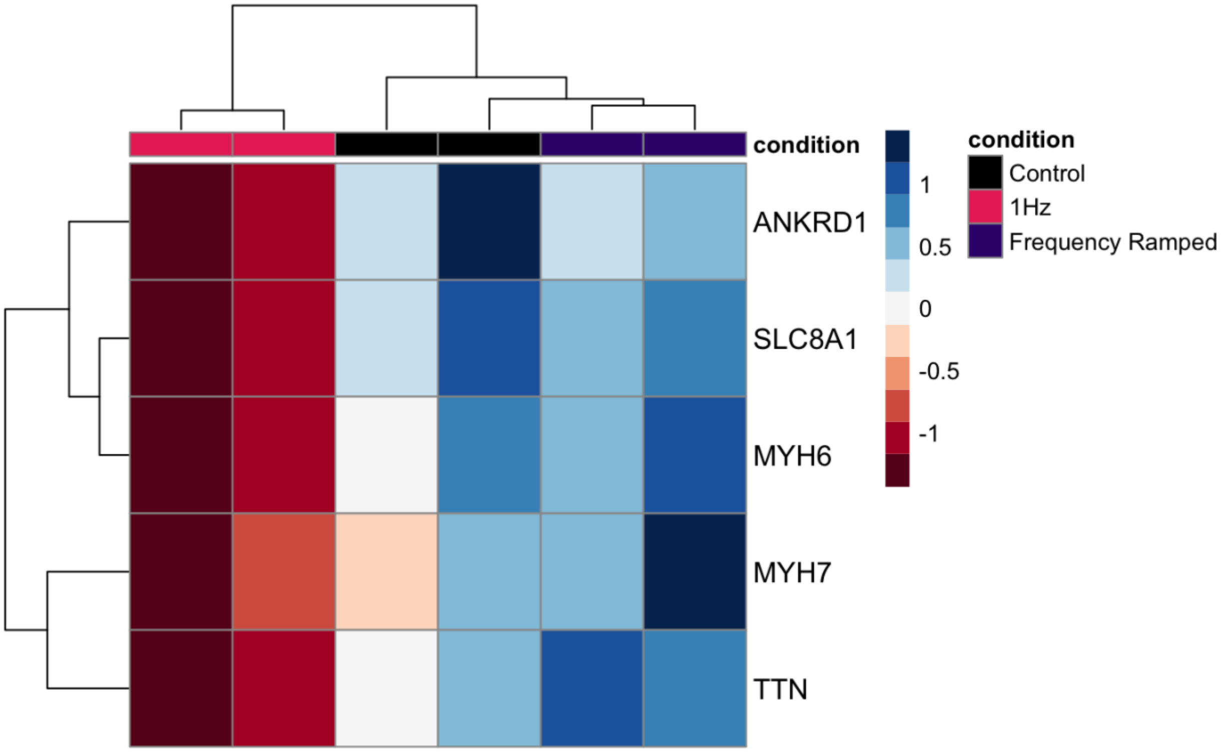


**S6: Highest Differentially Expressed Cardiac Genes:** Decomposing the genes found in GSEA cardiac pathways, the top 5 most differentially expressed genes, across all conditions, are identified. Color bar represents z-score normalized within each row.

**Supplementary Table S1**

| Gene Name | Primer Sequence |
|---|---|
| TBP | 5'-GCTGTTTAACTTCGCTTCCG-3' |
| | 5'-CAGCAACTTCCTCAATTCCTG-3' |
| RYR2 | 5'-CAGAGTTCGCACAGTAACAGT-3' |
| | 5'-CAGCCAATCTCCAGTCCATT-3' |
| GJA1 | 5'-ACTTGGCGTGACTTCACTAC-3' |
| | 5'-GTACTGACAGCCACACCTTC-3' |
| CACNA1C | 5'-CTTCGAGTACCTGATGTTCGTC-3' |
| | 5'-GTTGAGGATGTTCATGGCGAT-3' |
| TNNT2 | 5'-AGCATCTATAACTTGGAGGCAGAG-3' |
| | 5'-TGGAGACTTTCTGGTTATCGTTG-3' |
| NR2F2 | 5'-CGAGTACAGCTGCCTCAAG-3' |
| | 5'-CTTCCAAAGCACACTGAGACT-3' |
| IRX4 | 5'-ATGTCCTACCCGCAGTTTG-3' |
| | 5'-GTGCTCAGGGAGTTGGTG-3' |
